# Reliable hypotheses testing in animal social network analyses: global index, index of interactions and residual regression

**DOI:** 10.1101/2021.12.14.472534

**Authors:** Sebastian Sosa, Cristian Pasquaretta, Ivan Puga-Gonzalez, F Stephen Dobson, Vincent A Viblanc, William Hoppitt

## Abstract

Animal social network analyses (ASNA) have led to a foundational shift in our understanding of animal sociality that transcends the disciplinary boundaries of genetics, spatial movements, epidemiology, information transmission, evolution, species assemblages and conservation. However, some analytical protocols (i.e., permutation tests) used in ASNA have recently been called into question due to the unacceptable rates of false negatives (type I error) and false positives (type II error) they generate in statistical hypothesis testing. Here, we show that these rates are related to the way in which observation heterogeneity is accounted for in association indices. To solve this issue, we propose a method termed the “global index” (GI) that consists of computing the average of individual associations indices per unit of time. In addition, we developed an “index of interactions” (II) that allows the use of the GI approach for directed behaviours. Our simulations show that GI: 1) returns more reasonable rates of false negatives and positives, with or without observational biases in the collected data, 2) can be applied to both directed and undirected behaviours, 3) can be applied to focal sampling, scan sampling or “gambit of the group” data collection protocols, and 4) can be applied to first- and second-order social network measures. Finally, we provide a method to control for non-social biological confounding factors using linear regression residuals. By providing a reliable approach for a wide range of scenarios, we propose a novel methodology in ASNA with the aim of better understanding social interactions from a mechanistic, ecological and evolutionary perspective.

## INTRODUCTION

Over the past 50 years, graph theory has established itself as an important methodology in the study of natural or artificial interconnected systems (anthropology^1^, sociology^2^, economics ^3^, ecology^4^, ethology^5^, animals societies^6^) whether at a microscopic (e.g., proteomic^7^) or macroscopic (e.g., ecosystem^8^) scale. In regards to the study of animal sociality, novel methods (i.e., indices of associations and pre-network permutations) building upon traditional graph theory have allowed to address issues specific to this field (e.g., heterogeneity in sampling effort). Hence, animal social network analysis (ASNA) has become an important methodology for major advances in both theoretical and empirical research on animal sociality^4,9^, social ontogeny^10^, genetic mechanisms of sociality^11^, impact on fitness^12^, epidemiology^10^, animal culture^10^, and social structures (e.g., social structure diversity and evolution^13-16^). These achievements were made possible thanks to the establishment of analytical methods that were considered state-of-the-art protocols.

ASNA typically has two key-objectives: 1) estimating sociality patterns among individuals, quantified in a social network and 2) testing statistical hypotheses regarding patterns of sociality among individuals (as described further). To accomplish the first objective, researchers calculate a measure of the tendency to associate (i.e., undirected behaviour) or interact (i.e., directed behaviour)^17-19^ (i.e., social index) for each pair of individuals (dyad). The resulting set of values is used to construct a social network, with each individual as a node and the values of the social index providing an index of the strength of the connection (edge weight) between members of each dyad. To account for individual heterogeneity in sociality and observation frequency, Hubalek^17^, Sailer^18^ and Whitehead^19^ provided a theoretical background to measure social affiliations with association indices. This has become a fundamental building block for describing social structures in ecological research. Many of the association indices that exist^6^ estimate the proportion of time that a pair of individuals spends in association. The higher the value of the index, the greater the level of association within the dyad. The most frequently used association index to date is the simple ratio index -SRI- (*Eqn*. 1), designed for data that were collected in discrete sampling periods.

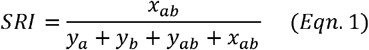

*x*_*ab*_ is the number of sampling periods with *a* and *b* observed associated, *y*_*a*_ is the number of sampling periods with only *a* identified, *y*_*b*_ is the number of sampling periods with only *b* identified and *y*_*ab*_ is the number of sampling periods with *a* and *b* identified but not associated.

However, association indices only provide a measure of associations between individuals, but do not inform on the direction of social interactions. In contrast, interaction networks are often constructed when a particular type of interaction is the focus. In contrast to associations, interactions are usually directional (e.g., preening in birds), and can differ from association networks because the directionality of interactions is considered (e.g., bird *A* preens bird *B* frequently, but bird *B* does not preen bird *A*) seldomly. In this paper, we focus initially on association networks, but discuss how the methods can be extended to interaction networks in the second part of the paper.

To accomplish the second objective (testing statistical hypotheses about sociality), node-based measures are often used. Node-based measures are calculated from the network and they measure aspects of each individual’s position in the network^20^ (e.g., individual frequencies of associations/interactions: *strength*). Hypotheses testing typically relates to node-based measures, such as whether males have a higher *strength* than females. However, node-based measures cannot be used in a straightforward manner to test hypotheses about individual associations, because associations (as well as social interactions) are statistically dependent: they are counted twice, once for each individual of the dyad. This dependence violates parametric test assumptions by artificially inflating the degrees of freedom. Because degrees of freedom represent the number of values in the final calculation of a statistic that are free to vary, their inflation can generate high rates of false positives^21^. In addition, associations may be influenced by sampling biases and non-social factors^22^. The consequence may be a biased conclusion about the significance of the individuals’ associations, when one focuses on traditional testing against a null hypothesis (H_0_). Consequently, much of the methodological work in ASNA has focused on developing techniques that enable valid hypotheses testing. Such techniques are needed to provide a low rate of false negatives, that is the non-rejection of a false hypothesis (e.g., a conclusion that males and females on average have similar *betweenness*, when in fact males have higher *betweenness*). At the same time, one must avoid the risk of inflating false positives, that is the rejection of the null hypothesis when it is true (e.g., a conclusion that males have a higher *strength* than females, when in fact there is little difference between the sexes). More specifically, we would want to reject a null hypothesis that is true at the 5% significance level only 5% of the time: any more than this is an inflation of the false positive error rate. The use of parametric statistics in social network analyses typically results in inflated error rates due to the problem of non-independence in the data described above.

One way to achieve valid hypothesis testing in ASNA is to resort to the use of permutation techniques^23^. Permutation methods consist first in computing a test statistic on real data (e.g., a *t* statistic for the difference in *strength* between males and females), then randomizing the original dataset a certain number of times (e.g., 10,000 times) while keeping any pattern that may have existed in the real data (e.g., males no longer have a higher *strength*), and finally the test statistic is recalculated from the permuted data to yield a null distribution for the test statistic, which is used in place of a parametric null distribution (e.g., a *t* distribution in our example) - this statistic is taken to be a representative value if the null hypothesis were correct-. In ASNA, permutation^23^ techniques can take two forms, network or pre-network permutations. A variety of network permutation approaches exist (see Hobson, et al. ^24^ for a review) and the most frequently used is the node label permutation approach that consists of randomizing individuals’ characteristics rather than aspects of their sociality. Instead of randomizing individuals’ characteristics, pre-network permutations (also called data stream permutations)^23^ are performed on the raw data, after each permutation the network is reconstructed and a new test statistic is generated for the null distribution. The manner of randomization must be carefully chosen, so that it disrupts the pattern being tested for (e.g., males having higher *strength* than females), but also maintains any statistical dependencies in the data that might cause a false positive. Consequently, pre-network permutations usually consist of generating a randomized alternative dataset constrained by the biological features of the original dataset by maintaining 1) the same total number of individuals, 2) the same number of dyads observed, 3) the same number of times each individual is observed and 4) the same number of individuals in each group. It must be noted, though, that pre-network permutations have been used primarily for the study of associations and not for the study of interactions. This is perhaps because interactions are often directed in nature, e.g., bird *A* attacks bird *B*, or chimpanzee *C* grooms chimpanzee *D*, and the definition of association-based indices like the SRI (*eq*. 1) only allow for undirected relationships. For the study of individual interactions, researchers have mainly applied network permutations that were directly performed on networks that were calculated using indices designed specifically for interaction data and allow the study of directed social behaviors (e.g., allogrooming, aggression) (see Croft, et al.^12^ for further details). Nonetheless, the combination of indices of associations with both pre-network permutations and network permutations have become standard testing protocols in ASNA, but have recently been called into question^25,26^.

Recent studies have identified important reliability issues related to the pre- and network permutation analytical protocols, such as inflated rates of false negatives (i.e., non-rejection of a false null hypothesis) and false positives (i.e., acceptance of a false null hypothesis) for both permutation approaches^25,26^. For example, Puga-Gonzalez, et al.^25^ used simulations for a case with data biases arising from the data collection protocol (e.g., oversampling of specific individual categories), and found that pre-network permutations showed rates of false positives of 35.6%, while network permutations showed rates of false negatives of 60.8% and rates of false positives of 36.6%. Yet, very few biological data collected *in natura* are immune to biases related to the system under study or related to necessarily limited sampling. Given the central role of pre-network and network permutations in ASNA, high rates of false negatives and false positives are likely to be a common problem and a pressing issue to resolve for ensuring the reliability of hypotheses testing in ASNA.

Whereas important statistical advances have helped improve the reliability of statistical hypothesis testing^20,22^ in ANSA, limitations remain. Franks, et al.^20^ proposed the use of linear regression for testing, while adding control factor(s) to account for potential biases. However, this approach was only validated for undirected association data using pre-network permutations and inclusion of additional variables in the model reduce degree of freedom. Farine & Carter 2021^22^ suggested an approach using both permutation processes to estimate the deviance from randomness, but this approach still returns high rates of false positives and can only be used for association data. Finally, Hart, et al. ^27^ recently demonstrated that parametric tests without permutations show rates of false positives and false negatives similar to those of network permutation tests, thereby calling into question the use of permutations themselves. Moreover, although authors argue that permutations do not control for data dependency, the purpose of permutations is to provide an alternative to compute a test statistic against null models and to avoid reducing the degree of freedom of parametric tests. As a result, it currently remains difficult to identify a standard analytical protocol in ASNA, according to the type of data collected and the data collection protocol.

Here, we propose an approach for addressing the issue of high rates of false positives and false negatives. The main idea behind our approach is that indices of associations measure the proportion of time that a dyad spends in association, however, while the computation of these indices considers the sampling effort, indices of associations are calculated using absolute time (i.e., a SRI of 0.5 means that a dyad spends 50% of the time associated, regardless of whether individuals were observed 10 times or 100 times). Thus, in order to account for sampling effects, indices of associations need to be weighted to obtain a value relative to the sampling effort. We term this the “global index” (GI) approach. In addition, as indices of associations have been designed only for undirected behaviours, we also developed an “index of interactions” (II) that estimates the proportion of interactions of a dyad, while accounting for both received and given behaviours and allowing the use of GI for directed behaviours.

Sampling biases are only one aspect of the problem and, as described by Farine & Carter 2021^22^, these factors can be of two different types. First, indices of associations may be altered by sampling biases (i.e., observation time heterogeneity) related to the data collection protocol or to individuals’ specificities (e.g., cryptic individuals are more challenging to observe). The second potential confounding factor that may influence individual associations refers to non-social biological influences of sociality such as cycle synchrony across individuals, space use or kinship, among others. The consideration of potential confounding non-social biological influences on sociality is of major interest in order to correctly evaluate the effect of sociality. While Farine & Carter 2021^22^ proposed some solutions to control for such confounding factors, these show the same limitations previously discussed (high rates of false positives and applicable only to association data). Moreover, developing a methodology allowing to reach beyond the control for putative biological confounding factors, by assessing their magnitude or importance on social interactions, remains to be done. In the third part of our study, we propose a method to control for non-social biological confounding influences (e.g., sex, age, body condition). Our approach uses linear regressions to estimate the relationship between an assumed non-social biological confounding factor and a social measure. If the relationship is significant, we can then consider that the non-social biological confounding factor affects the individual social measure. We can “control” for the factor by computing the residuals from the linear regression and using them as a relative social measure. This approach, defined as the “residual correction” (RC), has the advantage of being usable after accounting for sampling biases (after using GI), using permutation approaches to compute significant relationships, estimating whether one or several non-social biological confounding factor(s) exist, and statistically controlling for these factors. Furthermore, the use of generalized linear mixed models allows accounting for structure of the data (e.g., repeated measurements and non-Gaussian distributions like Poisson, or zero-inflated distributions).

Finally, it is possible to test for non-linear relationships between the social measure and the potential non-social biological confounding factor(s) through polynomial regressions.

We perform computer simulations and first demonstrate that the GI approach (that consists in considering individuals’ sampling effort within the indices of associations) is reliable for the study of undirected behaviors. In a second step, we show that the index of interactions (II) can be reliably used in combination with the GI to study directed behaviours. Finally, we show that the RC approach accurately estimates and controls for non-social biological confounding factors. Our simulations mimic a number of different sampling (focal sampling or scan sampling or “gambit of the group” -described below-) and recording protocols^29^ (continuous and timed sampling) commonly used to collect ASNA data, highlighting that our methods generate reliable results, there are observation biases, variations in the data collection protocol, or the type of behaviour studied (directed/interactions or undirected/associations).

## METHOD & RESULTS

### Issue related to control of observations - time heterogeneity

While indices of associations accurately estimate differences in associations among the individuals from different dyads, numerous confounding factors may affect these associations. Several studies have attempted to control for such confounding factors. For example, as highly gregarious individuals associate more frequently with other highly gregarious conspecifics just by chance, even if there is no mutual affiliation, Godde et al.^30^ suggested consideration of the individuals of a dyad. Similarly, in order to account for the specificities in gregariousness between categories of individuals, Peeper et al.^31^ suggested some modifications to the computation of association indices. The Peeper et al. 1999 and Godde et al. 2013 approaches are conceptually similar and modify the indices of associations as follows:

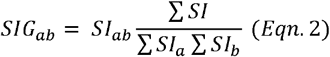

where *SI*_*ab*_ is the social index for the dyad *ab*, ∑ *SI* is the sum of all the SIs for all dyads, and ∑ *SI*_*a*_ and ∑ *SI*_*b*_ are the sums of all the SIs for individuals *a* and *b*. Note that the authors used a specific social index (the half weight index-a modification of the SRI), but the logic can be applied to any appropriate social index of association that measures the proportion of time spent associating. The logic is that if patterns of association were governed only by each individual’s levels of gregariousness, we would expect the proportion of time *a* and *b* spend in association to be proportional to ∑ *SI*_*a*_ ∑ *SI*_*b*_. This would represent the case where individuals associate with others at random. Consequently, dividing by ∑ *SI*_*a*_ ∑ *SI*_*b*_ “corrects” the social index, *SIG*_*ab*_, such that it quantifies the tendency of *a* and *b* to associate after accounting for their individual gregariousness. As ∑ *SI* is a constant for all dyads, it can be removed so we obtain:

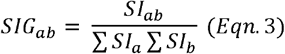

where the denominator controls for gregariousness of each individual of the dyad.

Here we propose a similar method (global index: GI, *Eqn*. 4) to control for differences in sampling time among individuals, by replacing ∑ *SI*_*a*_ ∑ *SI*_*b*_ with the sampling time of each individual of the dyad. This will weight each proportion of time that two individuals spend together by the inverse of each individual’s total time of observation:

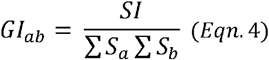

where ∑ *S*_*a*_ is the sampling effort of individual *a* and ∑ *S*_*b*_ is the sampling effort of individual *b*. Note that this formula can be only used with weighted social network measures and not with binary social network measures, such as the *degree* (i.e., number of social partners), that only consider the presence/absence of links without considering their weights. Therefore, for binary social network measures, we propose division of the social network measure by the sum of the sampling efforts of the individuals for whom the social network measure is computed and those of its social partners. Once the GI is used to construct the social network, the same index can be used as part of a pre-network permutation process in order to test hypotheses about the network. We hypothesize that this correction will solve many of the problems with sampling biases described above, and test our hypothesis below using computer simulations.

Concretely, if we consider discrete time sampling rule^29^ (instantaneous or one-zero sampling) such as spatial associations collected with “gambit of the group” sampling protocol (i.e. considering spatially clustered individuals associated), or behavioral events without duration collected with scan sampling protocol, sampling effort of an individual *a* would simply be the number of sampling periods in which it was observed. Similarly, if we consider continuous recording sampling rule of behavioral state with (e.g. time of grooming) or without (e.g. frequencies of grooming) duration collected with focal sampling protocol^29^, the sampling effort of an individual *a* would be the time of all focal bouts made on *a*. The two following simulations mimic both cases of behavioural recording: continuous behavioural frequencies collected through focal sampling and discrete behavioural sampling (e.g., spatial associations collected through GoG) to highlight how the GI performed under these sampling and recording rules for undirected behaviours using the simple ratio index (i.e., 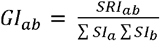).

### Simulations to validate the GI method for undirected behaviours

To demonstrate the reliability of the Global Index (GI) approach in multiple scenarios, we used the simulation from Puga-Gonzalez et al.^14^. This simulation, inspired by Farine^23^, simulates datasets collected by focal sampling in which individuals of opposite sexes show differences in sociality, whether statistically significant or not. In addition, it is possible to mimic a specific amount of sampling protocol bias by simulating oversampling for males. With such a simulated dataset, it is possible to test for differences in sociality between sexes through network permutations, pre-network permutations, and parametric tests. Hence, this simulation allows the assessment of rates of false negative error, while simulating a significant difference in sociality between males and females with or without the presence of sampling biases. On the other hand, when simulating similar sociality levels in males and females with or without the presence of sampling bias, it is possible to assess rates of false positive error. As in the simulations of Puga-Gonzalez et al. 2020^25^, we used Latin hypercube sampling^32^ with the ‘lhs’ R library^33^ to sample the parameter space (variables a–d in Table 1). 500 different combinations of input parameter values were run per simulation scenario, producing a total of 2,000 simulations.

**Table 1.**
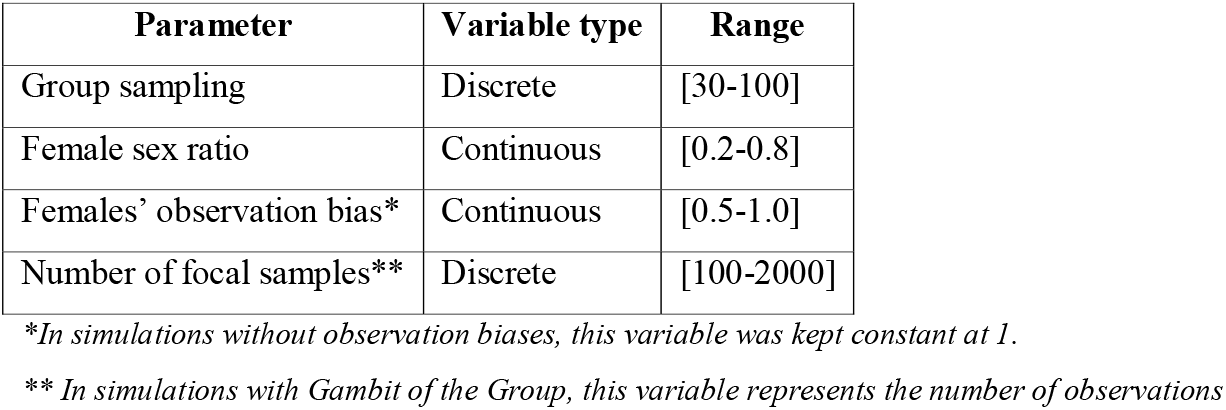
Range of variation of initial parameters for simulations 1, 2 and 3

| Parameter | Variable type | Range |
| --- | --- | --- |
| Group sampling | Discrete | [30-100] |
| Female sex ratio | Continuous | [0.2-0.8] |
| Females' observation bias* | Continuous | [0.5-1.0] |
| Number of focal samples** | Discrete | [100-2000] |

\*In simulations without observation biases, this variable was kept constant at 1.
\*\* In simulations with Gambit of the Group, this variable represents the number of observations

In addition, we made two modifications to the original simulation. The first consisted of using the GI approach. To compute the GI (see *Eqn*.*4*), we computed first the *SRI* (see *Eqn*.*1*). The second was to compute individuals’ *degree* and *eigenvector* to validate our method also with measures that consider individuals’ direct links only (first-order measures: *degree* and *strength*) and a measure that considers individuals’ indirect links (second-order measure: *eigenvector*)^20^ with the R package ANTs^34^. These changes were the only ones we made regarding the evaluation of GI reliability for focal sampling (simulation 1, Appendix 1). However, in order to evaluate the GI reliability for GoG, we made another modification by considering each observation as a cluster of associated individuals and not a focal observation (simulation 2, Appendix 2).

Results for simulation 1 (focal sampling) showed important improvement for network permutations using the GI approach for focal sampling with rates of false negatives below 5% with or without the presence of observation biases and rates of false positives below 5% with observation biases and below 10% without observation biases for all social network measures (Table 2). As expected, parametric tests showed high rates of false positives related to over-inflation of the degree of freedom used to calculate the significance^35^. Finally, the GI approach did not solve the issue related to pre-network permutations (i.e., it did not address the null hypothesis that *X* was distributed randomly with respect to *Y* or that the effect of *X* on *Y* was zero) and thus, as expected, the GI did not solve the issue of high rates of false positives using the pre-network approach.

**Table 2.** Results for simulation 1. Overview of global index (GI) percentage of false positive/negatives for undirected behaviours with focal sampling data collection protocol according to scenarios with and without observation bias and different hypothesis testing approaches.

|  | Simulation with biases of observation |  |  |  |  |  | Simulation without biases of observation |  |  |  |  |  |
| --- | --- | --- | --- | --- | --- | --- | --- | --- | --- | --- | --- | --- |
|  | False negative rates (%) |  |  | False positive rates (%) |  |  | False negative rates (%) |  |  | False positive rates (%) |  |  |
|  | Degree | Strength | Eigenvector | Degree | Strength | Eigenvector | Degree | Strength | Eigenvector | Degree | Strength | Eigenvector |
| Parametric test | 5 | 2.6 | 0.6 | 6.6 | 40.2 | 30 | 2.4 | 0.4 | 0 | 3.2 | 3.6 | 3.6 |
| Network permutation test | 4.6 | 1 | 0.4 | 3.4 | 0.6 | 0.2 | 0 | 0.4 | 0 | 3.6 | 3.4 | 4 |
| Pre-network permutation test | 65.8 | 16.4 | 28.4 | 38 | 13.2 | 23 | 42.2 | 65.8 | 59.2 | 60 | 15.2 | 14.2 |

Results for simulation 2 (scan sampling/GoG) showed similar values for reliability (Table 3) with rates of false positives below 5% with or without the presence of observation biases for all social network measures. Finally, whereas rates of false negatives were under 5% with observation biases for all social network measures, these rates reached 93.4% in simulation without biases and for the *degree* measure. This issue derives from the fact that most of simulations generated fully-connected networks in which all individuals had an equal number of partners (equal to the defined group size) and without biases of observation. While we could have modified the simulation algorithm to avoid the creation of fully-connected networks, scan/GoG sampling protocols often generate dense networks and, therefore, we chose to keep the simulations as such as a warning regarding the use of binary network measures under scan/GoG sampling protocol.

**Table 3.** Results for simulation 2. Overview of global index (GI) percentage of false positive/negatives for undirected behaviours with GoG data collection protocol according to scenarios with and without observation bias and different hypothesis testing approaches.

|  | Simulation with biases of observation |  |  |  |  |  | Simulation without biases of observation |  |  |  |  |  |
| --- | --- | --- | --- | --- | --- | --- | --- | --- | --- | --- | --- | --- |
|  | False negative rates (%) |  |  | False positive rates (%) |  |  | False negative rates (%) |  |  | False positive rates (%) |  |  |
|  | Degree | Strength | Eigenvector | Degree | Strength | Eigenvector | Degree | Strength | Eigenvector | Degree | Strength | Eigenvector |
| Parametric test | 1.2 | 0 | 0 | 98.6 | 14.4 | 13 | 97 | 0 | 0 | 5.6 | 6.8 | 6.8 |
| Network permutation test | 1 | 0 | 0 | 98.8 | 2 | 2.4 | 96.8 | 0 | 0 | 4.8 | 5.2 | 5.2 |
| Pre-network permutation test | 100 | 100 | 100 | 0 | 0 | 0 | 100 | 100 | 100 | 0 | 0 | 0 |

**Table 4.** Results for simulation 3. Overview of global index (GI) percentage of false positive/negatives for directed behaviours with focal sampling data collection protocol according to scenarios with and without observation bias and different hypothesis testing approaches.

|  | Simulation with biases of observation |  |  |  |  |  | Simulation without biases of observation |  |  |  |  |  |
| --- | --- | --- | --- | --- | --- | --- | --- | --- | --- | --- | --- | --- |
|  | False negative rates (%) |  |  | False positive rates (%) |  |  | False negative rates (%) |  |  | False positive rates (%) |  |  |
|  | Outdegree | Outstrength | Outeigenvector | Outdegree | Outstrength | Outeigenvector | Outdegree | Outstrength | Outeigenvector | Outdegree | Outstrength | Outeigenvector |
| Parametric test | 2.2 | 5.4 | 0.2 | 4.6 | 6.2 | 6.8 | 1.4 | 6.2 | 0.4 | 6.2 | 4.2 | 6 |
| Network permutation test | 1.6 | 3.4 | 0 | 4.2 | 4.8 | 6.6 | 1.6 | 5.4 | 0.2 | 6.8 | 4.6 | 7.8 |

### Extension of GI approach to directed behaviours

While individuals’ associations are mostly used in behavioral ecology in which entire populations are followed over large areas, the study of individuals’ social interactions are also an important part of ASNA research. The study of social interactions is usually done in smaller and well-identified groups. In order to enable reliable testing for questions about social interactions, we developed an index of interactions (II) and evaluated the reliability of the combination of II with GI through a third simulation.

The appropriate form of an index of social interactions (the II approach) for directed behaviours depends on the recording rule used to collect data, which depends in part on the nature of the target interaction. Here we showed that for interaction data collected in discrete sampling periods (instantaneous or one-zero sampling), a modified version of the SRI is appropriate, but that if the interaction data were collected using continuous recording, then a simple ratio is more appropriate.

The first possibility is that the target interaction is a behavioural state of a meaningful duration, e.g., bouts of grooming. The researcher might then wish to estimate the proportion of time that each dyad (*ab*, consisting of individuals *a* and *b*) spends engaged in the target interaction. Under such cases, instantaneous sampling may be used^29^, with the data specifying whether the target interaction was occurring for each dyad at uniform time points (e.g., every 5 mins). In this case the II approach is directly analogous to the collection of standard association data, therefore the SRI (Eqn. 1) can be used except, *x*_*ab*_ is the number of sampling points at which *a* and *b* were observed interacting with one another, *y*_*a*_ is the number of sampling points at which only *a* was identified, *y*_*b*_ is the number of sampling points at which only *b* was identified and *y*_*ab*_ is the number of sampling points at which *a* and *b* were identified but not engaged together in the target interaction. Again *y*_*null*_, here the number of points at which neither *a* nor *b* were identified, was not included in the calculation. In simple terms, the II approach calculated the proportion of points at which *a* and *b* were observed interacting, but excluded the cases where the researcher is not sure if they were interacting or not.

It is also straightforward to extend the II approach to directed interactions, e.g. if we want to separately estimate the proportion of time *a* spends grooming *b*, and the proportion of time *b* spends grooming *a*.

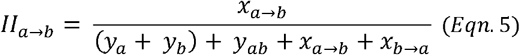

Here *x*_*a*→*b*_ is the number of sampling points at which *a* was directing an interaction towards *b* (e.g., *a* was grooming *b*) and *x*_*b*→*a*_ is the number of sampling points at which *b* was directing an interaction towards *a* (e.g., *a* was grooming *b*). Therefore, for such data the SRI, or the II approach (Eqn. 2), can be used under the same assumptions as for association data: e.g., failing to observe individuals *a* and *b* while they are interacting is as likely as failing to observe them both when they are not interacting together (see Hoppitt & Farine 2018).

Alternatively, the target interaction may be a behavioural event of short duration (e.g., one bird directs a peck at another bird), or the researcher may simply be interested in the rate at which *a* initiated interactions with *b*, rather than the proportion of time engaged in that interaction. In such cases one-zero sampling is traditionally used^29^. Here the data specify whether or not at least one event *a* → *b* occurred in each sampling period of length *X*. Here we could intuitively use the II approach given in Eqn. 5, with suitably modified definitions of the terms: e.g., *x*_*a*→*b*_ is now the number of sampling periods during which *a* was observed directing an interaction towards *b, y*_*a*_ is the number of sampling periods for which only *a* was identified and *y*_*ab*_ is the number of sampling periods for which *a* and *b* were identified but not engaged together in the target interaction. Here the directed *II*_*a*→*b*_ estimates the probability of seeing at least one interaction directed from *a* to *b* in a time period of length *X*, again under equivalent assumptions as the SRI for association data^36^. The researcher may prefer to convert this to an estimate of the rate at which interactions *a* → *b* occur as *λ*_*a*→*b*_ = −*log*(1 − *II*_*a*→*b*_)/*X*.

### Simulation on directed behaviours: Focal sampling and continuous recording

In order to simulate datasets of directed interactions collected through focal sampling with continuous recording (simulation 3, Appendix 3), we modified simulation 1 by assigning directionality to what was originally considered associations. For simplicity, we considered all observations as emitted behaviours. For each simulated dataset, we computed the GI (see *Eqn.4*) using the II (see *Eqn.5*) as social index. This allowed us to assess the reliability of the II and GI, with or without sampling biases and with or without relationship between emitted behaviours and individuals’ sex. We ran node label permutations and parametric tests using the Latin hypercube sampling with the ‘lhs’ R library^33^ to sample input parameter space (Table 1). For each scenario, we ran 500 different combinations of input parameter values for a total of 2,000 simulations. In this simulation, as well as in the following one, we did not perform pre-network permutations for two reasons. First, because data stream permutations for directed behaviours do not exist, and second because the GI approach does not solve pre-network permutation reliability issues related to hypothesis testing, as discussed earlier.

Results of simulation showed that the combination of II and GI returns low rates of false positives and false negatives, with or without sampling biases for node label permutations and parametric test (Table 3). The low rates of false positives and false negatives of parametric tests in these simulations for directed behaviours are supported by the fact that, as these data are independent (an emitted interaction is counted once), over-inflation of the degree of freedom should not occur even when testing hypothesis in scenarios with observational biases. This is why, in such scenarios, parametric tests should be preferred. However, it is quite common that researchers use several directed social network measures (total emitted behaviours and total received behaviours) in a linear regression to evaluate their effect. In such cases, data independence is violated as an individual’s emitted behaviours are the received behaviours of its congeners and thus a single behaviour is counted twice. In this scenario, network permutations should be preferred.

### Simulation on directed behaviours: Scan sampling and discrete time sampling

In order to simulate datasets of directed interactions collected through discrete time sampling, we created a simulation that mimics scan sampling (simulation 4, Appendix 4). This model creates a population of size *N* with a predefined number of subgroups (*g*, 4 subgroups in each simulation) within the population. By creating subgroups, we created *cliques* with groups of individuals having higher probability of being observed together, yet seldomly interacting, thereby shaping the *y*_*ab*_ term of the II in Eqn.5. A number of scans (*x)* was predefined at initialization of the simulation. For each observation, a within and between subgroups interaction process was defined. In the within subgroup interaction process, a subgroup (*g*) was selected randomly, a number of individuals (*n*) observed within the scan (*i)* was defined following a Poisson distribution (with alpha of 6). Once the size of the scan was defined, *n*/2 individuals belonging to the subgroup *g* were selected, and these individuals (defined as *emitters*) will emit an interaction based on a fixed probability (*P*_1_) towards a selected subgroup member. In this way, and according to the probability *P*_1_, we can create a network with a given amount of preferential attachment (i.e. individuals interact preferentially or not with the same individuals). In the case where an *emitter* did not have social partners, an interaction toward a social partner belonging to subgroup (*g)* was randomly created. In the between groups interaction process, a fixed probability (*P*_2_) was defined to enable within the scan (*i)* the presence of individual(s) that do not belong to the selected subgroup *g*. For each of these individuals, an incoming link from an individual belonging to the selected subgroup *g* was created. This interaction processes (within and among subgroups) were iteratively repeated until the defined set of observations (*x)* reached a desired value. The simulation input parameters (N, x, *P*_1_, *P*_2_) are listed in Table 5.

**Table 5.**
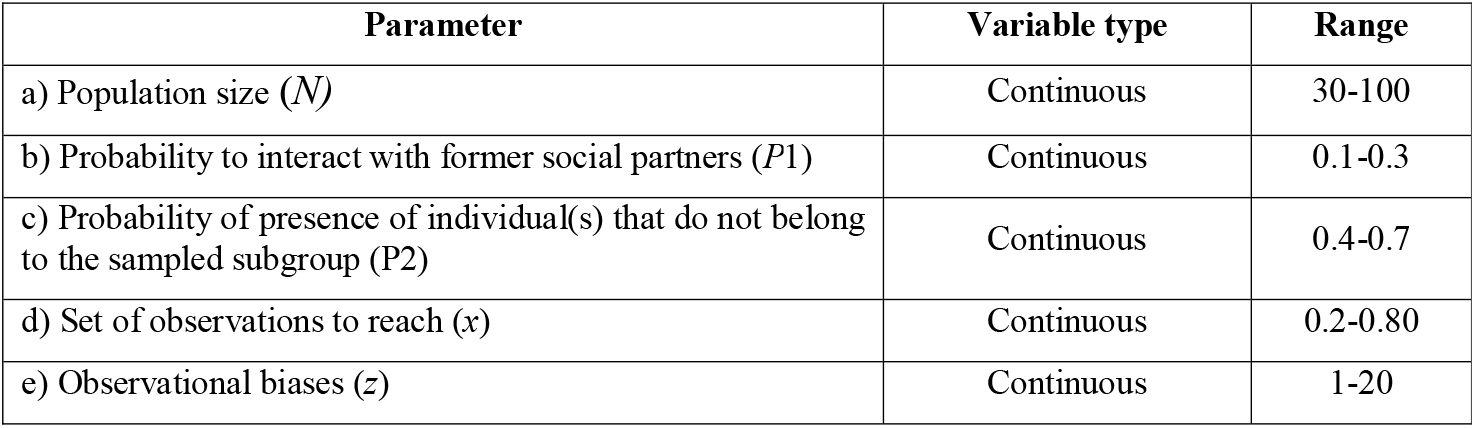
Range of variation of initial parameters for simulation 4

Once the simulation was done, we calculated the GI (see Eqn.4) using the II (see Eqn.5) as social index and the directed versions of the previous social measures were computed for each individual through II and GI: 1) *outstrength, outdgree* and *outeigenvector*. To assess the reliability of hypothesis testing, a random continuous variable (*x*) following a normal distribution was created. This variable, that represents an individual trait, was then ordered and assigned based on individuals’ social measures, to create a relationship between *x* and social measures (scenario 1), and also randomly assigned to individuals’ strength to create no relationship between *x* and social measures (scenario 2).

In order to mimic observational biases (*z*) proportional to the relationship between the variable *x* and the social measures, a maximum observational bias was defined (e.g. 20%) as input parameter of the simulations (Table 5), and this percentage decreased proportionally to the relative position of the individual according to its value *x*. For example, the individual showing the highest *x* had its number of observations reduced by 20%. The second individual (the one with the second highest *x*) had its number of observations reduced by 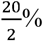, the third individual by 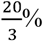, and so on. This allowed us to create scenarios with or without biases in combination with scenarios 1 and 2 explained above.

To evaluate the testing reliability of II combined with the GI approach, we performed 500 simulations for scenario 1 and 500 simulations for scenario 2. This was done with and without observational biases for a total of 2,000 simulations. We sampled the input parameter space of the simulations (variables a–e in Table 5) using the Latin hypercube sampling using ‘lhs’ R library. For each simulation, we assessed the rates of false negatives and false positives of network permutations and parametric tests.

Results of simulation showed that the combination of II and GI returned low rates of false positives and false negatives, with or without sampling biases for node label permutations and parametric test (Table 6). When observation biases were simulated, we observed that parametric test still returned no false positive nor false negative results, whereas we started to observe some false negative and false positive results with network permutations, although the rates still fell within acceptable levels (Table 6).

**Table 6.** Results for simulation 4. Overview of global index (GI) percentage of false positive/negatives for directed behaviours with scan sampling data collection protocol according to scenarios with and without observation biases and different hypothesis testing approaches.

|  | Simulation with biases of observation |  |  |  |  |  | Simulation without biases of observation |  |  |  |  |  |
| --- | --- | --- | --- | --- | --- | --- | --- | --- | --- | --- | --- | --- |
|  | False negative rates (%) |  |  | False positive rates (%) |  |  | False negative rates (%) |  |  | False positive rates (%) |  |  |
|  | Outdeg<br>ree | Outstren<br>gth | Outeigenvec<br>tor | Outdeg<br>ree | Outstren<br>gth | Outeigenvec<br>tor | Outde<br>gree | Outstre<br>ngth | Outeigenve<br>ctor | Outde<br>gree | Outstreng<br>th | Outeigenvect<br>or |
| Parametric test | 0.6 | 0 | 8.4 | 2 | 0 | 1.8 | 3.6 | 0 | 0.2 | 2.6 | 0 | 2.2 |
| Network permutation test | 0.4 | 3 | 8.4 | 2 | 0.6 | 1.8 | 3.6 | 0 | 0.2 | 2.4 | 0.2 | 2.2 |

**Table 7.**
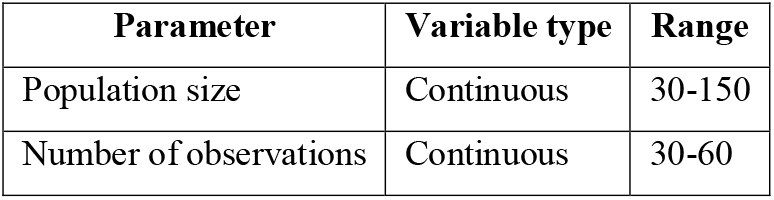
Range of variation of initial parameters for simulation 5

**Table 8.** Results for simulation 5. Overview of regression correction (RC) combined with global index (GI) percentage of false positives for undirected behaviours with GoG data collection protocol according to scenarios with non-social biological confounding factors.

|  | Without RC (%) |  |  | With RC (%) |  |  |
| --- | --- | --- | --- | --- | --- | --- |
|  | Degree | Strength | Eigenvector | Degree | Strength | Eigenvector |
| *Parametric test* | 84.06 | 99.9 | 100 | 5.6 | 0.15 | 3.06 |
| *Network permutation test* | 87.52 | 100 | 100 | 5.53 | 0.59 | 8.16 |

### Estimating and controlling for non-social biological confounding factors

Finally, to be able to estimate and control for non-social biological confounding factors, we used residuals of the regression between the social measure and the potential non-social biological confounding factors. As residuals represent the difference between the prediction of the linear regression model of the relationship between the predictive variable (the potential non-social biological confounding factor(s)) and the predicted variable (the social measure), they allowed us to adjust the social measure according to the potential non-social biological confounding factor(s), and use it as relative social measure. As discussed in the introduction, the residual correction (RC) method has the advantage of being usable after accounting for sampling biases (after using GI), of using permutation approaches to compute significant relationships, and of estimating whether one or several non-social biological confounding factor(s) exist. Furthermore, it is possible to resort to generalized linear mixed models in order to account for structure of the data (e.g., repeated measurements, non-Gaussian distribution such as Poisson, or zero-inflated distributions). Finally, non-linear relationships can be tested between the social measure and the potential non-social biological confounding factor(s) with the use of polynomial regressions.

To assess the reliability of RC, we used the Farine & Carter 2021^22^ simulation (simulation 5, Appendix 5) that mimics association data collected on a population using GoG sampling and a discrete time recording rule. Each individual had a trait value *T*_*i*_ drawn from a normal distribution. By assigning individuals with the highest trait values to the largest observed groups *X* (ranging from 1 to 10) or by assigning individuals to observed groups randomly, the simulation created, respectively, scenarios where individuals’ traits impacted their spatial associations or scenarios where individuals’ traits had no impact on their spatial associations. In addition, a conceptual modification to the simulation was done to mimic the effect of a non-social biological confounding factor. This modification consisted of considering individuals’ group size *X* as the result of spatial preference, where individuals in large groups live on large patches that contain more resources and therefore have a greater carrying capacity. With this modification, individuals’ spatial associations were no longer considered as a result of individual social decisions only, but rather as an outcome of habitat heterogeneity and individual space use. This is an ideal simulation to evaluate the reliability of the RC approach as in such scenario the social measure is related to individuals’ traits but individuals’ sociality is an indirect outcome of individuals’ spatial preferences. In addition, Farine & Carter 2021^22^ simulation showed that standard permutation tests (pre-network and network permutations) return high rates of false positives while this should not occur because this social measure is an outcome of habitat heterogeneity and individual space use as discussed previously.

We ran 39,600 of the modified simulations sampling the parameter space with 100 simulations for each possible combination of population size (ranging from 30 to 150) and number of observations per individuals (ranging from 30 to 60). For each simulated dataset, we computed the *strength, eigenvector* and *degree* and performed analyses with and without the RC approach for node label permutations. We expect to observe no significant relationship between individuals’ traits and spatial associations when using RC, whereas a significant relationship should be found without RC (as highlighted by simulations in Farine & Carter 2021^22^). Thus, the RC approach accurately estimates and controls for non-social biological confounding factors for both first- and second-order network measures. As in simulation 1, rates of false positives and false negatives with or without observation biases for the *degree* were very high for the same reasons as explained in simulation 1 (i.e., GoG simulation creates highly connected networks). Finally, when the RC approach was not used, the rates of false positives increased drastically, indicating that it is important to control for non-social biological confounding factors in order to obtain reliable statistical results.

## DISCUSSION

In the present study, we developed a Global Index (GI), an approach that weights dyadic association/interaction indices according to their respective sampling effort. Simulations show that this method returns acceptable rates of false negatives and false positives errors, with or without biases of observations. The GI approach can be used for both directed and undirected behaviours using focal sampling, scan sampling or Gambit of the Group data collection protocols, and can be used for first-order (*degree* and *strength*) and second-order (*eigenvector*) social network measures. Our simulations show that pre-network permutations as well as parametric tests for ASNA return unacceptable rates of false negatives and false positives, even using the GI approach, and suggest these should be avoided for ASNA research to ensure reliable hypothesis testing. Finally, we also provide a method to estimate and control for non-social biological confounding factors using the residuals of individuals’ social measure values regressed on the estimate of confounding factors, showing reliable results. One major asset of this approach is that it can be combined with the GI to account for multiple confounding factors at once and takes into account the data structure (e.g., repeated measurements, spatial or phylogenetical observation clustering, non-Gaussian distribution). However, while the original simulations from Ivan et al. 2020 and Farine & Carter 2021 simulated datasets with some sample sizes lower than 30, we modified the minimum sample size (N ≥ 30) for all simulations, as these use parametric tests (linear regressions) to test statistical hypotheses. Further tests might determine whether GI and II, in combination with non-parametric tests, provide reliable statistical results.

Together with the growing interest and use of graph theory for research on social complexity, variance analysis (e.g., intraclass correlation coefficient for the study of repeatability) is starting to be used in ASNA^37^ and, to date, hypothesis testing reliability for those approaches have not been tested and should thus be considered cautiously. Similarly, temporal analyses of individuals’ sociality is an important part of ASNA to understand sociality dynamics arising from demographic^38,39^ variation, environment^40^, and ontogeny^41^. However, as for variance analyses, those require further testing of the reliability of the mixed models that are used to study them. Nonetheless, our results show that high rates of false negatives and false positives are not related to the permutations themselves but rather to an issue with control of observation time heterogeneity. We expect that “node label” permutations with GI approach we propose here is also reliable for variance analysis or other more complex hypothesis testing approaches, although further tests are needed. By providing a reliable approach for a wide range of scenarios, we propose a novel methodology in ASNA with the aim of better understanding animal sociality and animals’ societies from a mechanistic, ecological and evolutionary perspective.

## AKNOWLEDGMENT

This research was supported by fellowship grant from the Institute of Advanced Studies of the University of Strasbourg to F.S.D. and V.A.V. F.S.D. thanks the Région Grand Est and the Eurométropole de Strasbourg for the award of a Gutenberg Excellence Chair. We also thank Benjamin Beltzung for the discussions regarding index of sociality issues.

